# Imaging cortical engagement during motor imagery, mental arithmetic, and silent word generation using MEG beta rhythm

**DOI:** 10.1101/2022.07.16.500303

**Authors:** Vahab Youssofzadeh, Sujit Roy, Anirban Chowdhury, Aqil Izadysadr, Lauri Parkkonen, Manoj Raghavan, Girijesh Prasad

**Affiliations:** Department of Neurology, Medical College of Wisconsin, Milwaukee, USA; BrainAlive Research Pvt Ltd, India; School of Computer Science and Electronic Engineering, University of Essex, UK; Department of Psychology and Criminal Justice at Middle Georgia State University, USA; Department of Neuroscience and Biomedical Engineering, Aalto University School of Science, Espoo, Finland; Intelligent Systems Research Centre, School of Computing, Engineering, and Intelligent Systems, Ulster University, Derry ∼ Londonderry, N. Ireland, UK

**Keywords:** Magnetoencephalography (MEG), Beta-decrements, Brain-computer interface (BCI), Task imagery, Classification

## Abstract

Accurate mapping of cortical engagement during mental imagery or cognitive tasks remains a challenging brain-imaging problem with immediate relevance to the development of brain-computer interfaces (BCI). We analyzed data from fourteen individuals who performed cued motor imagery, mental arithmetic, or silent word generation tasks during MEG recordings. During the motor imagery task, participants imagined the movement of either both hands (HANDS) or both feet (FEET) after the appearance of a static visual cue. During the cognitive task, participants either mentally subtracted two numbers that were displayed on the screen (SUB) or generated words starting with a letter cue that was presented (WORD). The MEG recordings were denoised using a combination of spatiotemporal filtering, the elimination of noisy epochs, and ICA decomposition. Cortical source power in the beta-band (17–25 Hz) was estimated from the selected temporal windows using a frequency-resolved beamforming method applied to the sensor-level MEG signals. The task-related cortical engagement was inferred from beta power decrements within non-overlapping 400 ms temporal windows between 400 and 2800 ms after cue presentation relative to a baseline 400 ms temporal window before cue onset. We estimated the significance of these power changes within each parcel of the Desikan-Killiany atlas using a non-parametric permutation test at the group level. During the HANDS and FEET movement-imagery conditions, beta-power decreased in premotor and motor areas, consistent with a robust engagement of these cortical regions during motor imagery. During WORD and SUB tasks, beta-power decrements signaling cortical engagement were lateralized to left hemispheric brain areas that are expected to engage in language and arithmetic processing within the temporal (superior temporal gyrus), parietal (supramarginal gyrus), and (inferior) frontal regions. A leave-one-subject-out cross-validation using a support vector machine (SVM) applied to beta power decrements across brain parcels yielded accuracy rates of 74% and 68% respectively, for classifying motor-imagery (HANDS-vs-FEET) and cognitive (WORD-vs-SUB) tasks. From the motor-versus-nonmotor contrasts, accuracy rates of 85% and 80% respectively, were observed for HAND-vs-WORD and HAND-vs-SUB. A multivariate Gaussian process classifier (GPC) provided an accuracy rate of 60% for a four-way (HANDS-FEET-WORD-SUB) classification problem. The regions identified by both SVM and GPC classification weight maps were largely consistent with the source modeling findings. Within-subject correlations of beta-decrements during the two task sessions provided insights into the level of engagement by individual subjects and showed moderately high correlations for most subjects. Our results show that it is possible to map the dynamics of cortical engagement during mental processes in the absence of dynamic sensory stimuli or overt behavioral outputs using task-related beta-power decrements. The ability to do so with the high spatiotemporal resolution afforded by MEG could potentially help better characterize the physiological basis of motor or cognitive impairments in neurological disorders and guide strategies for neurorehabilitation.

## 1. Introduction

Tasks designed for the development of brain-computer interfaces (BCI) often involve mental simulation of actions without their actual execution (Szameitat et al., 2007; Crammond, 1997; Jeannerod, 1994). In contrast to mapping brain responses driven by external sensorimotor stimuli, the mapping of brain areas activated by such tasks remains a key challenge in BCI research (Wolpaw et al., 2000), both due to the distributed nature of brain networks that are engaged during cognitive tasks and the limitations of different imaging methods. The high spatial and temporal resolution of magnetoencephalography (MEG) makes it inherently suitable for mapping the dynamical engagement of brain areas during mental processes, and thus, recent interest in MEG in the context of BCI research (Parkkonen, 2015; Mellinger et al., 2007; Lai et al., 2005).

Specific aspects of task-induced changes in cerebral electrical activity are increasingly recognized as signatures of local cortical engagement during information processing in the brain (Giraud and Poeppel, 2012; Hauk et al., 2017; Lewis and Bastiaansen, 2015; Meyer, 2018; McFarland et al., 2000; Salmelin and Hari, 1994; Schnitzler et al., 1997; Pfurtscheller and Neuper, 1997; Engel and Fries, 2010; Spitzer and Haegens, 2017). Induced oscillatory responses are time-locked, but not necessarily phase-locked to stimuli, and therefore cannot be extracted by averaging the time-domain cerebral responses, but they may be detected as power changes in different frequency bands (Pfurtscheller and Andrew, 1999; David et al., 2006; Başar and Bullock, 1992; Tallon-Baudry, 1999). Over the last few decades, a number of studies using EEG, Electrocorticography (ECoG), and MEG have demonstrated increases in gamma band (> 40 Hz) power concurrent with a decrease in power in the alpha (8-12 Hz) and beta (13-30 Hz) bands as a characteristic feature of the cortical response to afferent stimuli (Crone et al., 2006, 1998; Miller et al., 2007; Eulitz et al., 1996; Pfurtscheller, 1991; Singh et al., 2002; Wagner et al., 2012). High-Gamma activity appears to be related to the firing rates of local neuronal populations (Manning et al., 2009; Nir et al., 2007; Ray and Maunsell, 2011; Edwards et al., 2005; Michalareas et al., 2016) and is more focally expressed than concomitant decreases in alpha/mu and beta power (Crone et al., 2006, 1998; Miller et al., 2007; Eulitz et al., 1996; Pfurtscheller, 1991; Singh et al., 2002; Wagner et al., 2012). The high spatiotemporal resolution of MEG is ideally suited to capturing such time-frequency dynamics and localizing their cortical sources (Gross, 2019).

While gamma-band activity arising outside primary sensory or motor cortices may be less readily detectable in M/EEG (e.g., compared to ECoG) task-related power decrements in the beta-band consistent with the expected cortical engagement have been demonstrated by several studies (Youssofzadeh et al., 2020; Weiss and Mueller, 2012; Seeber et al., 2014; Neuper and Pfurtscheller, 2001). Importantly, the decrease in beta-band power has been observed in motor imagery (Pfurtscheller et al., 2006; Klepp et al., 2015) as well as during various cognitive tasks (Weiss and Mueller, 2012; Armeni et al., 2019; Lewis and Bastiaansen, 2015). For example, beta power decreases have been found to be correlated with imaginary foot movements when walking on a virtual street (Pfurtscheller et al., 2006; Neuper and Pfurtscheller, 2001). A pioneering EEG-BCI study showed that feature values based on beta-band activity over the sensorimotor area provided the largest discrimination, > 90% and 80% for execution and imagination, respectively (Bai et al., 2008). On the other hand, the suppression of beta during language processing has been associated with novel or unexpected stimuli (Engel and Fries, 2010; Weiss and Mueller, 2012; Bastiaansen et al., 2010), semantically incongruous sentences (Luo et al., 2010; Wang et al., 2012), and unexpected high-versus-low perplexity (Armeni et al., 2019). Many M/EEG studies support the utility of monitoring beta power in the localization of language functions in healthy controls and patients (Findlay et al., 2012; Fisher et al., 2008; Grabner et al., 2007; Kadis et al., 2008; Passaro et al., 2011; Weiss and Mueller, 2012; Youssofzadeh et al., 2020).

In this study, we estimated task-induced beta-band (17–25 Hz) power decrements in the cortex from MEG recordings performed during movement-imagery involving the hands or feet, mental arithmetic, and silent word generation in a group of healthy individuals (N = 14). We demonstrate that source-space task-related beta-band power decrements can map cortical engagement during mental processes that are not attributable to dynamic sensory stimuli or overt behavioral responses. This is supported by reasonably high classification accuracy and consistent ROIs suggested by the classification weight maps. For ease of reporting, we use the term “beta-decrements” to refer to decreased source-power in the beta-band frequency relative to the pre-cue baseline period (aka, event-related desynchronization, or ERD, or suppression effects) throughout the paper.

## 2. Material and Methods

### 2.1. Participants

We analyzed MEG data from 14 participants who performed cued motor imagery, mental arithmetic, or word generation during MEG recording. Demographic details of the participants have been reported previously (Rathee et al., 2021). They were recruited from the community through advertising or were students and staff of the University of Ulster; all had normal hearing and normal or corrected-to-normal vision. Individuals with a history of neurological or psychiatric illness and individuals taking psychoactive medication were excluded. The participants were 11 men and 3 women, with a mean age of 29.3 years *±* 5.9 (SD), 12 right-handed, and 2 left-handed, based on the self-reported questionnaire. The participants signed a written consent form before commencing the experiments. The ethics committee at the University of Ulster, Northern Ireland, UK, had approved the experimental protocol.

### 2.2. Data acquisition

MEG data were recorded using a 306-channel (204 planar gradiometers and 102 magnetometers) whole-head neuromagnetometer system (Elekta Neuromag TRIUX; MEGIN Oy, Helsinki, Finland) in the upright position in a magnetically shielded room (ETS-Lindgren, Eura, Finland) located at the University of Ulster, Magee campus, Northern Ireland, UK. The raw data were acquired at a sampling rate of 1 kHz and high-pass filtered with a cutoff frequency of 0.03 Hz. The cues for the tasks were displayed on a projector screen (a Panasonic projector with a screen resolution of 1024×768 pixels and a refresh rate of 60 Hz). Participants were seated in a comfortable chair approximately 80 cm away from the projector screen. Further details can be found in an earlier paper (Rathee et al., 2021; Roy et al., 2020).

### 2.3. MEG task paradigm

Participants completed two sessions of a BCI experiment during MEG recording. Sessions were recorded on different days for each participant. The paradigm required the completion of four mental tasks when the appropriate cue was presented visually: imagining the movements of both hands (HANDS), the movements of both feet (FEET), mental-subtraction of two digits (SUB), and the generation of a word starting with the cued letter (WORD), as illustrated in Fig. 1. Each task trial started with a pre-cue period lasting 2 s, then the appearance of a static visual cue, which remained on the screen for 5 s while the participant performed the cued task, followed by 1.5–2 s of rest. During the pre-cue period, a red fixation cross (“+”) was presented along with an auditory tone (500 Hz). During the cued-task period, a visual cue was presented corresponding to each of the 4 tasks (HANDS, FEET, WORD, or SUB). During the HANDS condition, the cue was a picture of two hands, and participants imagined grasping with both hands. During the FEET condition, the cure was a picture of two feet, and participants imagined dorsiflexing both feet. During the SUB condition, a subtraction problem (the subtraction of a 2-digit from a 3-digit number) was displayed on the screen for the participant to mentally execute. During the WORD condition, participants mentally generated a word starting with the letter cue shown on the screen, e.g., BOY for the letter B. The letters were randomly selected from the alphabet, A-Z (N = 26). For each task condition, 50 trials were acquired for a total of 400 trials (2 × 50 × 4, corresponding to the session, trial, and task conditions) per subject. The order of the tasks was randomized within each session.

**Figure 1:**
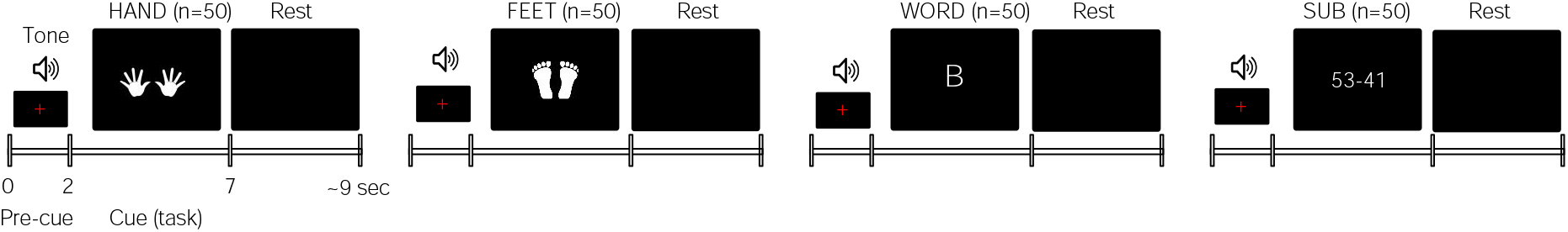
BCI task paradigm. Participants took part in 2 sessions of a task-imagery (BCI) MEG experiment. The task consisted of two motor- (Hands and Feet movements) and two cognitive (Word generation and mathematical Subtraction) tasks. A red fixation cross (“+”) and a tone were presented during the pre-cue period. During a cue-task period, a picture was presented corresponding to each of the 4 imagery task conditions of HANDS, FEET, WORD, and SUB. During the HANDS condition, participants imagined grasping with hand movement. During the FOOT, participants imagined movement of the foot upward. During the WORD, participants generated a word starting with a letter shown on the screen. The letters were randomly selected from A-Z. During the SUB condition, participants completed a 2-digit against 2-digit subtraction task.

### 2.4. Data analysis

We analyzed the MEG responses following cue onset (i.e., the period corresponding to task performance) using seven non-overlapping 400-ms temporal windows (from 400 to 2800 ms). The initial 400 ms after cue onset was excluded in order to discard transient sensory responses to the cue-onset. We compared responses during these temporal windows to a 400-ms period immediately preceding cue-onset, which served as a baseline (see Section 2.5 for details).

#### 2.4.1. Data preprocessing

Using MaxFilter software ver. 2.2 (MEGIN Oy, Helsinki, Finland), a temporal variant of signal space separation was applied to remove external magnetic interference and interpolate the signals of noisy sensors (Taulu and Simola, 2006). Data were epoched from –1 to 4 s relative to the cue onset, then bandpass-filtered (Butterworth with an order of 4) in a frequency range of 1 to 40 Hz. Segments containing artifacts (signal jumps, eye blinks, head movements, or muscle contractions) were removed by a threshold value defined by a variance exceeding 3 *·* 10^*−*24^ T, a kurtosis larger than 15, and a z-score larger than 4. Cardiac artifacts were removed via independent component analysis using the Infomax algorithm (Bell and Sejnowski, 1995). An average (*±* SD) of 8 *±* 6 trials per session were omitted.

A time-frequency response (TFR) analysis of sensor-level data was conducted to inspect the presence of beta-power changes. The TFR analysis was conducted using multitapers in the range of 1–50 Hz. Using sensors, a frequency-dependent sliding time window was analyzed in a time and frequency range of -400 ms to 3 s, and a three-cycle-long Hanning window (ΔT= 3/f, f is the frequency of interest) was used. Fourier representation was estimated using a spectral smoothing of ΔF =0.8x f. The TFRs were baseline-corrected based on the 400-ms pre-cue data. A sample TFR from an individual completing a HANDS imagery task is shown in Fig. 2.

**Figure 2:**
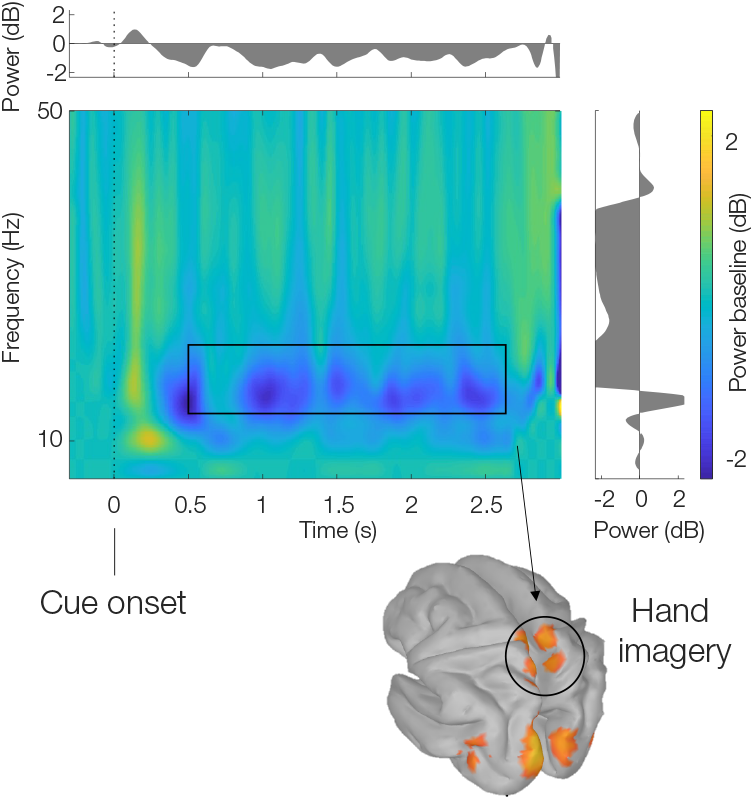
Time-frequency responses of a hand imagery task. An average of all sensor-level event-related power changes relative to baseline (−300, 0 ms) was utilized to support the selection of beta-band frequency, as specified by a rectangle. For illustrative purposes, corresponding beta-band power deceased source activities from an individual completing a hand movement motor imagery task are shown.

### 2.5. Frequency beamforming source analysis

The MEG data were coregistered to a T1-weighed anatomical MRI template (ICBM152). Cortical surface reconstruction was performed on cortical pial surfaces, downsampled to 15002 vertices. Overlapping spheres were used as a head model to estimate the lead-field matrix. Beta-band source power was estimated using Dynamic Imaging of Coherent Sources (DICS), a frequency-resolved spatial filtering beamforming technique (Gross et al., 2001). DICS uses the data covariance matrix to calculate the spatial filter based on the sensor-level cross-spectral densities (CSD), and the filter is then applied to the sensor-level CSD to reconstruct the source-level CSDs of the pairwise voxel activations. This provides coherent measures between the source pairs (off-diagonal elements of CSD) and the source power measures (diagonal elements of CSD). Our analysis only used the source power estimates.

Sensor-level CSD data were estimated in a beta-band frequency range of 17–25 Hz, using a center frequency of 21 Hz and a spectral smoothing window of 4 Hz. The Fourier transform and multitapering, with multiple tapers from Discrete Prolate Spheroidal Sequences, were used to estimate the CSDs (Slepian and Pollak, 1961). Beta-band source power during the task-performance period was contrasted against the 400-ms pre-cue baseline. To avoid biases due to unequal data segments, post-cue data were analyzed using 7 non-overlapping 400-ms time windows, ranging from 400 ms (to control for sensory responses) to 2800 ms. The average of source intervals was used as the representative beta power effects in each session. For mapping purposes, the average of the two sessions was used to represent the task conditions for each individual.Individual source findings were evaluated session-wise using machine-learning and correlation analyses.

### 2.6. Group source analysis

To achieve a group-level source analysis, a non-parametric permutation test was conducted (Maris and Oostenveld, 2007). Individual-specific power source values were projected to a default surface anatomy (MNI-152, nist.mni.mcgill.ca). An independent sample t-test was conducted against a null hypothesis to derive the t-statistics of each task condition. A Monte Carlo permutation test was applied with 5000 randomizations of extreme statistics. The extreme (maximum) statistics control the expected proportion of false positives (aka, multiple comparisons). A critical alpha of 0.05 was applied to the permutation distribution to report the significant statistical effects.

A Desikan-Killany (DK) atlas, consisting of 68 cortical regions (34 specific areas in L&R hemispheres) was used to summarize the power source measures (t-values) across regions (Desikan et al., 2006). The color-coded atlas regions are shown in Fig. 3. Regions of interests (ROIs) with corrected *t*-values with *p* < 0.05 were reported as being crucial to the task. Following the approach suggested by prior MEG studies (Youssofzadeh and Babajaniferemi, 2019; Papanicolaou et al., 2004; Tanaka et al., 2013; Raghavan et al., 2017), a conventional laterality index, LI = (L-R)/(L+R), was computed for the *t*-value of the left (L) and the right hemispheric (R) parcels to characterize the hemispheric involvement. LIs greater than 0.1 were considered left-lateralized, those less than -0.1 were considered right-lateralized, and intermediate ones were considered symmetric or bilateral.

**Figure 3:**
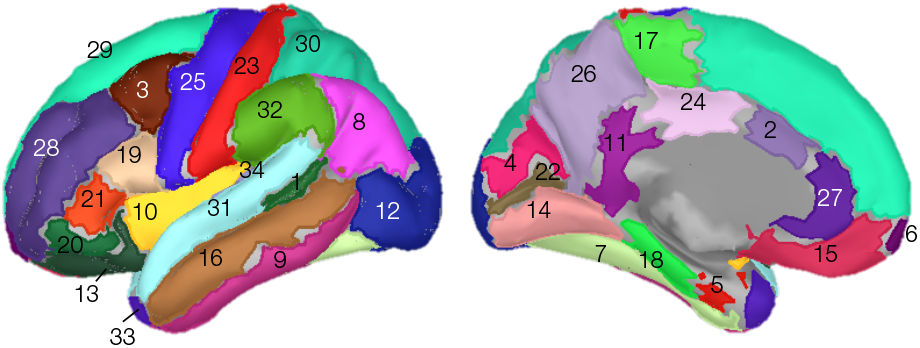
Desikan-Killany surface atla s. A Desikan-Killany atlas consisting of 68 (34×2) cortical regions distributed with Freesurfer (surfer.nmr.mgh.harvard.edu) was used to summarize source and pattern classification analyses. The regions were 1.bankssts; 2.caudalanteriorcingulate, 3.caudalmiddlefrontal; 4.cuneus; 5.entorhinal; 6.frontalpole; 7.fusiform; 8.inferiorparietal; 9.inferiortemporal; 10.insula; 11.isthmuscingulate; 12.lateraloccipital; 13.lateralorbitofrontal; 14.lingual; 15.medialorbitofrontal; 16.middletemporal; 17.paracentral; 18.parahippocampal; 19.parsopercularis; 20.parsorbitalis; 21.parstriangularis; 22.pericalcarine; 23.postcentral; 24.posteriorcingulate; 25.precentral; 26.precuneus; 27.rostralanteriorcingulate; 28.rostralmiddlefrontal; 29.superiorfrontal; 30.superiorparietal; 31.superiortemporal; 32.supramarginal; 33.temporalpole; 34.transversetemporal. Regions are randomly color-coded.

### 2.7. Pattern classification analysis

A pattern classification analysis was conducted to assess the discriminability of beta-decrements associated with the different tasks. Linear kernels per task condition were extracted as feature values from beta-decrements across the cortex. Linear kernels are pairwise similarity measures (dot product) between task source activations that are summarized in a kernel matrix (*N* × *N* dimensions, *N* : 14×2 subjects and sessions) (LaConte et al., 2005; Schrouff et al., 2013). To achieve an unbiased classification, kernels were mean-centered and normalized (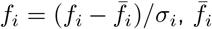 and *σ*_*i*_ are the mean and standard deviation of *i*-th feature, respectively). A linear support vector machine (SVM) with a default penalty parameter of C = 1.0 was applied to solve binary classification problems of HANDS *vs* FEET and WORD *vs* SUBtraction (Cortes and Vapnik, 1995). The SVM classifier relies on the assumption that two classes are separable by a linear decision boundary (separating hyperplane) in a feature space. In addition, a Gaussian process classifier (GPC) was used to classify all 4 BCI task conditions. The GPC is a probabilistic classification method relying on random field theory (Rasmussen and Williams, 2006), and has been successfully tested for the decoding of fMRI data(Marquand et al., 2010). For classification accuracy, class and balanced accuracy (BA, an average of sensitivity and specificity) were reported (Schrouff et al., 2013). The accuracy of the SVM and GPC models was evaluated by a leave-one-participant-out cross-validation procedure, and p-values were generated based on permutation testing with 1000 iterations. The DK surface atlas was used to summarize ROIs (regions of interest) from generated SVM and GPC weight maps. The weight maps are the spatial representation of the decision function and define the level of voxel contributions to the classification process. To summarize ROI contributions, the voxel weights were averaged within the defined anatomical regions by taking the sum of their absolute values and dividing them by the size of the region. We report classification results and weight maps using combined two-session beta-decrements. Our initial examinations based on permutation analysis suggested no significant classification rates for single-session data, likely due to the low sample size.

### 2.8. Correlation analysis

To assess the reproducibility of task-related cortical engagement in individual subjects, we performed two types of correlation analyses. We first asked how consistently, from session to session, does a subject’s task-related activation conform to the average map for the group for any particular task? To address this, we examined the correlation between beta-decrement maps for each task and the class-mean beta-decrement maps for the two sessions. The correlation was measured using a non-parametric linear bivariate Spearman test. The class mean was defined as the average beta-decrement map for all subjects across both sessions. These class-mean beta-decrements served as spatial templates for each task condition, against which the individual subject’s map was compared. Second, we ask, how reproducible is a subject’s activation map from session to session for any given task? To address this, we computed the direct between-session correlations of beta-decrements across the brain for each subject and task condition. We examined both absolute correlations (session 2 vs. 1) and their ratios (maximum/minimum). We hypothesized that a higher correlation to the class means and a higher correlation of activations across sessions indicate a higher level of engagement with the tasks. We also hypothesized that the consistency of the correlation between a subject’s activation map and the group mean across sessions may also identify the tasks for which activation maps are more stable.

## 3. Results

### 3.1. Group-level source analysis of beta-decrements

Source-level MEG activity during HANDS and the FEET motor imagery tasks showed significant beta-decrements in several cortical areas, including the precentral (the supplementary motor area, or SMA), the post-central, and the anterior cingulate gyri (Fig. 4.A). Based on the laterality-index, both tasks showed symmetric activation across the cerebral hemispheres, with LI_HANDS_ = 0.02 and LI_FEET_ = 0.04 (|LI|< 0.1). Source-level MEG activity during both the WORD and SUB task conditions showed significant beta-decrements in the temporal (superior temporal), parietal (supramarginal), and (inferior) frontal regions (Fig. 4.B). The laterality indices indicate a left-hemispheric dominance for both these tasks, with LI_WORD_ = 0.30 and LI_SUB_ = 0.26 (LI > 0.1).

**Figure 4:**
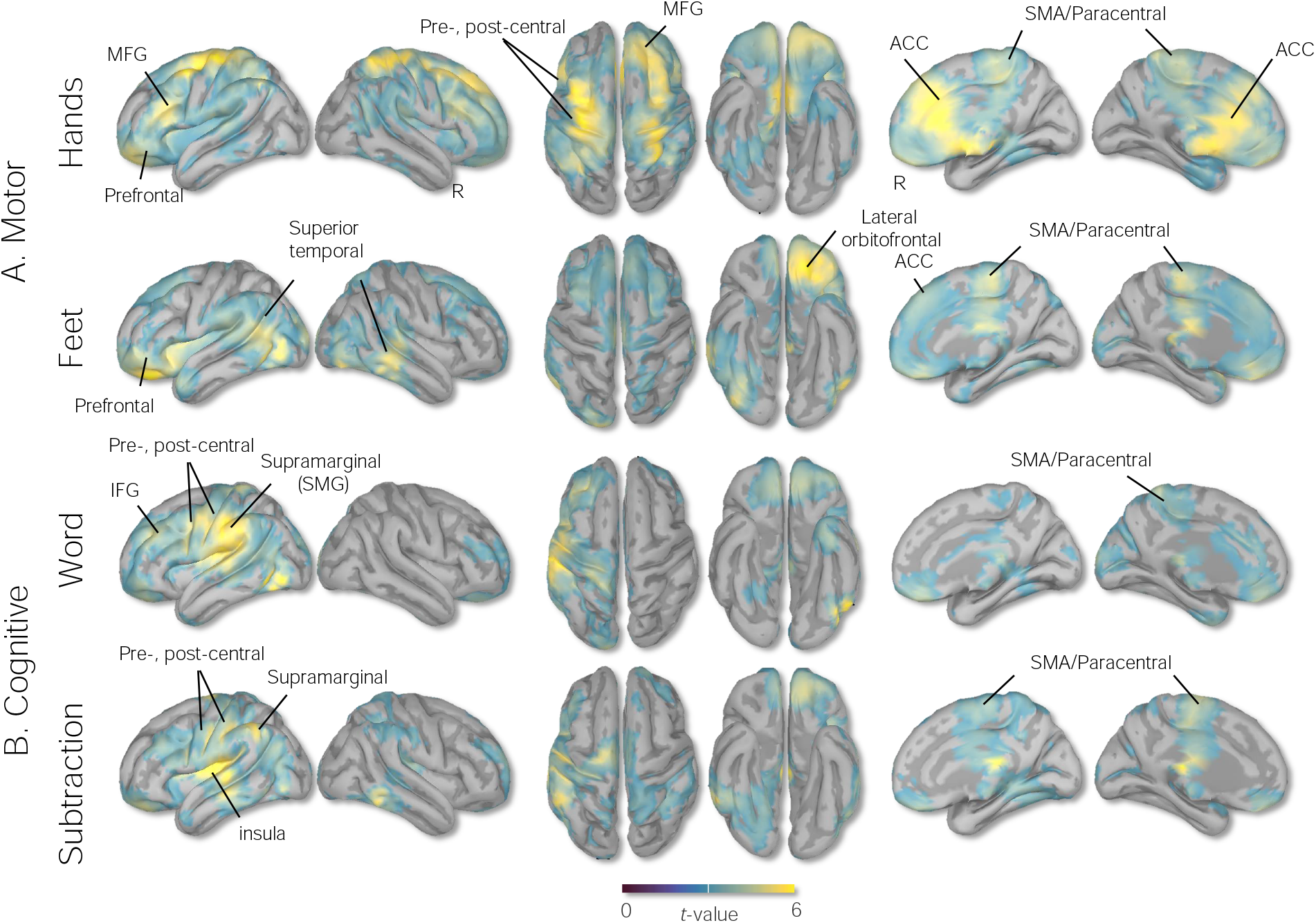
Group source analysis of task-imagery BCI experiment. Activation maps corresponding to (**A**) motor, Hands and Feet, motor imagery, and (**B**) cognitive, Word-generation, and Subtraction imagery tasks measured by the DICS beamformer source analysis in a frequency range of beta (17-25Hz) from 14 participants. *t*-maps are thresholded at a whole-cortex corrected *p* < 0.05.

The parcellation analysis of the HANDS task showed prominent cortical engagements bilaterally in the caudal anterior cingulate, middle frontal gyri, and paracentral regions. The FEET task showed cortical activations bilaterally in the paracentral (SMA), lateral orbitofrontal, banks of the superior temporal sulcus, and left precentral regions (Fig. 5A). The WORD task showed prominent left-hemispheric cortical engagements in the supramarginal, postcentral, precentral, and inferior frontal gyri (IFG) and the parsopercularis, while the SUB task revealed left-hemispheric cortical engagements within the temporal (transverse temporal and insula), parietal (supramarginal), and prefrontal (IFG, parsopercularis) regions (Fig. 5B, second row). The ROIs are summarized in Table 1.

**Figure 5:**
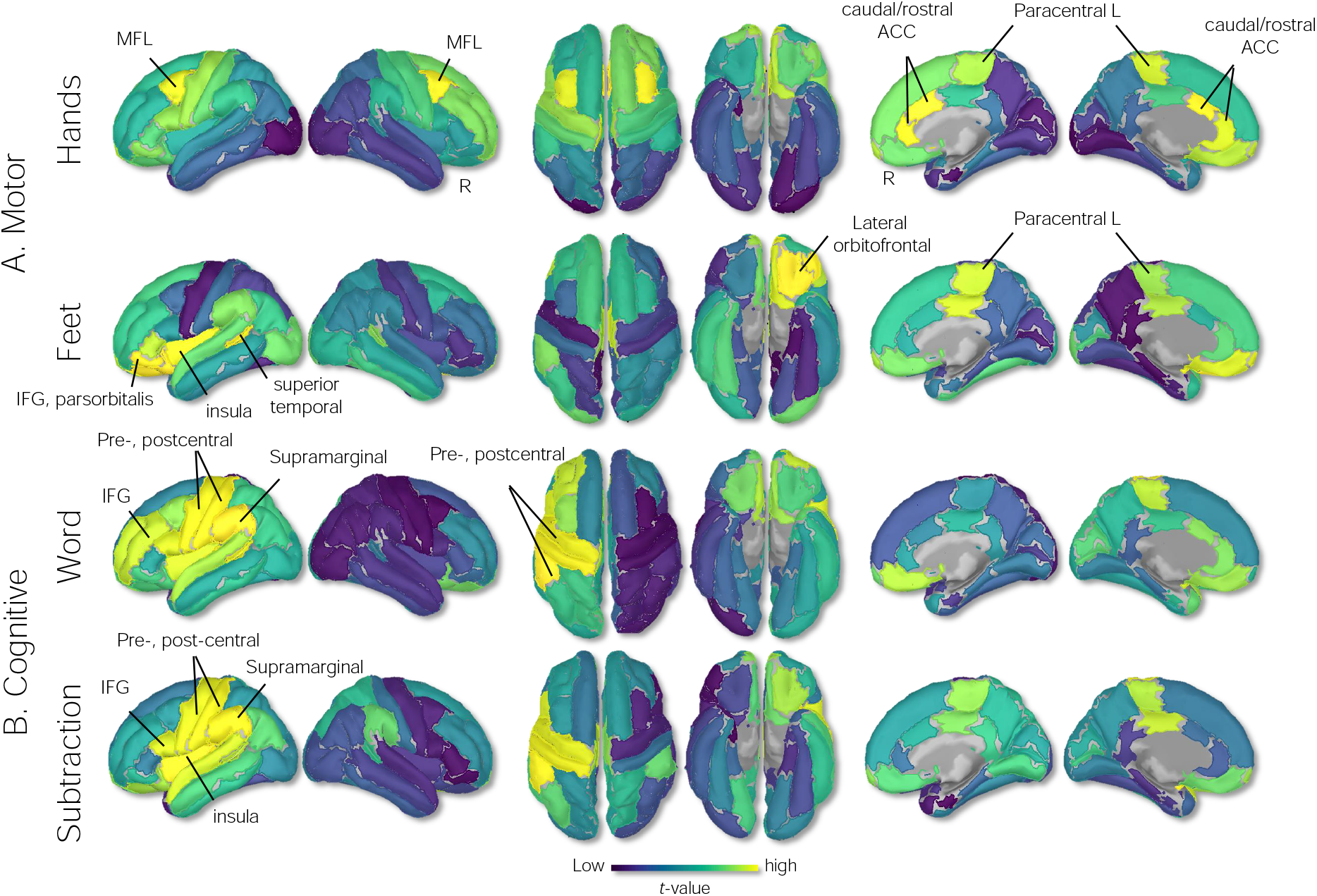
Group source parcellation analysis of task-imagery BCI experiment. Parcellation was conducted using a Desikan-killinay atlas (Fig.3). Yellow and blue color-coded areas represent higher and lower beta-decrements, respectively, consistent with t-values in Fig.4

**Table 1:** ROI involved in 4 task imagery BCI activities, as measured by beta-decrements.

| HANDS, ROI | t-value | FEET, ROI | t-value | WORD, ROI | t-value | SUB, ROI | t-value |
| --- | --- | --- | --- | --- | --- | --- | --- |
| caudal anterior cingulate R | 5.25 | lateral orbitofrontal L | 5.06 | supramarginal L | 5.04 | transverse temporal L | 5.54 |
| caudal anterior cingulate L | 4.81 | parsorbitalis L | 4.89 | transversetemporal L | 4.96 | insula L | 4.48 |
| caudal middle frontal R | 4.71 | bankssts L | 4.12 | postcentral L | 4.58 | supramarginal L | 4.38 |
| rostral anterior cingulate R | 4.69 | insula L | 4.11 | precentral L | 4.18 | postcentral L | 4.3 |
| caudal middle frontal L | 4.56 | frontal pole L | 4.05 | bankssts L | 3.92 | precentral L | 3.81 |
| rostral anterior cingulate L | 4.49 | medial orbitofrontal L | 4.03 | parsopercularis L | 3.82 | parsopercularis L | 3.77 |
| paracentral L | 4.39 | posteriorcingulate R | 3.96 | insula L | 3.8 | superiortemporal L | 3.67 |
| frontalpole L | 4.37 | paracentral R | 3.88 | rostral middlefrontal L | 3.74 | posteriorcingulate L | 3.67 |

### 3.2. Pattern classification analysis

The leave-one-subject-out cross-validation SVM pattern classification analysis (of beta-decrement maps) provided balanced accuracy (BA) rates of 74% and 68% for discriminating between the two motor-imagery tasks (HANDS-vs-FEET) and the two cognitive tasks (WORD-vs-SUB) tasks, respectively. Among all possible binary classification contrasts, the highest (and most significant) accuracy rates were achieved for the HANDS-vs-WORD and HANDS-vs-SUB classifications with BAs of 85% and 80%, respectively. The poorest (and non-significant) classification accuracy was achieved for the FEET-vs-SUB contrast, with a BA of 55%. A Gaussian process classifier (GPC) provided a BA of 60.36% for the four-way classification problem of HANDS-FEET-WORD-SUB. For ease of comparison, BAs are summarized in Table 2.

**Table 2:** BCI task classification accuracy using SVM and GPC classifiers. Summary classification accuracy of 7 different contrasts of BCI imagery tasks. Classification analyses were conducted using SVM and GPC classifiers and using linear kernels as input features. To ensure statistical independence, a leave-one-participant-out (LOO) cross-validation procedure with p-values generated using permutation testing with 1000 iterations was used. Class accuracy and balanced accuracy (BA, average sensitivity and specificity of class accuracy) are reported.

| Contrast | Total BA (%) | Class accuracy (%) | Method |
| --- | --- | --- | --- |
| HAND vs FEET | 74 (p = 0.04) | [72, 76] | SVM-LOO, Perm |
| WORD vs SUB | 68 (p = 0.05) | [72, 65] | SVM-LOO, Perm |
| HAND vs WORD | 85 (p = 0.007) | [85, 85] | SVM-LOO, Perm |
| HAND vs SUB | 80 (p = 0.02) | [76, 78] | SVM-LOO, Perm |
| FEET vs SUB | 55 (p = 0.2) | [60, 50] | SVM-LOO, Perm |
| FEET vs WORD | 75 (p = 0.04) | [77, 72] | SVM-LOO, Perm |
| HAND, FEET, WORD, SUB | 60 | [61, 64, 60, 54] | GPC-LOO |

The SVM weight maps generated for the HANDS *vs* FEET classification problem showed greater contributions from paracentral regions bilaterally (SMA) for the FEET condition, and greater bilateral contributions from the central regions (right postcentral and left precentral) and rostral anterior cingulate for the HANDS task. Also, the whole-cortex SVM weight maps generated for the WORD *vs* SUB classification problem revealed contributions from the left frontotemporal, and cingulate cortices (temporal lobe, anterior cingulate, supramarginal cortex, and IFG pars opercularis), and right temporal and parietal cortical regions (transverse temporal, inferior temporal, middle temporal, and superior parietal) for the SUB task conditions. Weight maps are shown in Fig. 6 and ROI contributions in weight percentage are summarized in Table 3. Consistent with SVM findings, the GPC weight maps showed high contributions by bilateral (pre-and post-) central gyri and the anterior cingulate gyri for the HANDS task, bilateral paracentral gyri for the FEET task, left temporal, parietal, and inferior frontal regions for the WORD task, and bilateral temporal and right parietal supramarginal region for the SUB task. Weight maps are shown in Fig. 7 and ROIs are reported in Table. 4.

**Figure 6:**
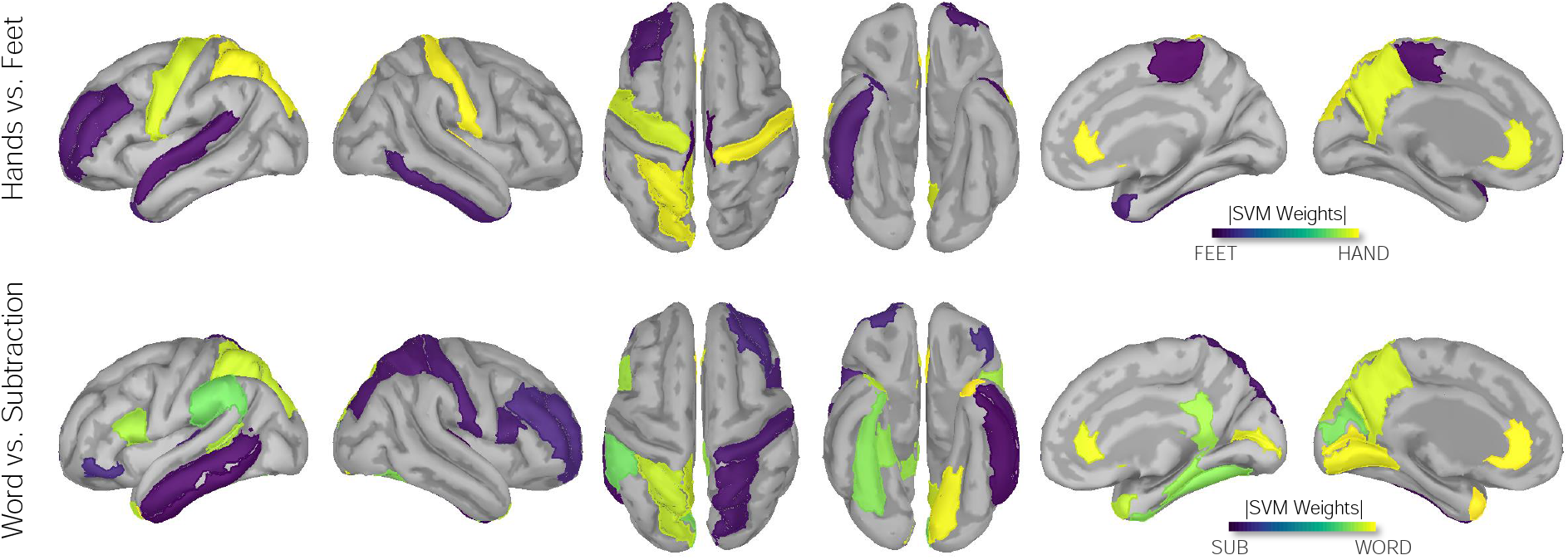
Weights (per region, Dk atlas) modeled by the SVM classification of BCI tasks. SVM classification weight maps of HANDS vs. FEET and WORD vs. SUB tasks are shown. Yellow and blue areas indicate higher SVM weights (i.e., greater beta-power source values) towards each side of the contrast. Maps are consistent with regions reported in Table. 3. Rendered weights are displayed on a surface template (MNI-152) with dark representing sulci and gray representing gyri. Suprathresholded values with values greater than half maximum on each side of contrast are shown.

**Table 3:** ROI involved in SVM classification weight maps of BCI tasks. Regions contributed (in percent) to the SVM classification of HANDS vs. FEET and WORD vs. SUB imagery tasks are reported. ROIs with percentage values greater than half-maximum (arbitrary 50% threshold) of each side of contrast are reported.

| **HANDS *vs* Feet, ROI** | **Weight (%)** | **Word *vs* Subtraction, ROI** | **Weight (%)** |
| --- | --- | --- | --- |
| transverse temporal R | 6.38 | temporal pole L | 5.7 |
| rostral anterior cingulate R | 5.01 | rostral anterior cingulate L | 5.69 |
| postcentral R | 4.91 | pericalcarine L | 5.35 |
| superior parietal L | 4.48 | lingual L | 5.13 |
| rostral anterior cingulate L | 4.37 | pericalcarine R | 4.83 |
| precentral L | 4.26 | rostral anterior cingulate L | 4.71 |
| posterior cingulate L | 4.19 | supramarginal L | 4.31 |
| postcentral | 4.15 | pars opercularis L | 4.22 |
| paracentral R | -5.93 | transverse temporal R | -5.1 |
| paracentral L | -5.4 | inferior temporal L | -5.0 |
| superior temporal L | -4.62 | middle temporal L | -4.62 |
| rostral middle frontal L | -4.6 | superior parietal R | -4.4 |
| inferior temporal R | -4.54 | postcentral R | -4.24 |
| temporal pole R | -4.45 | pars opercularis R | -4.01 |

**Figure 7:**
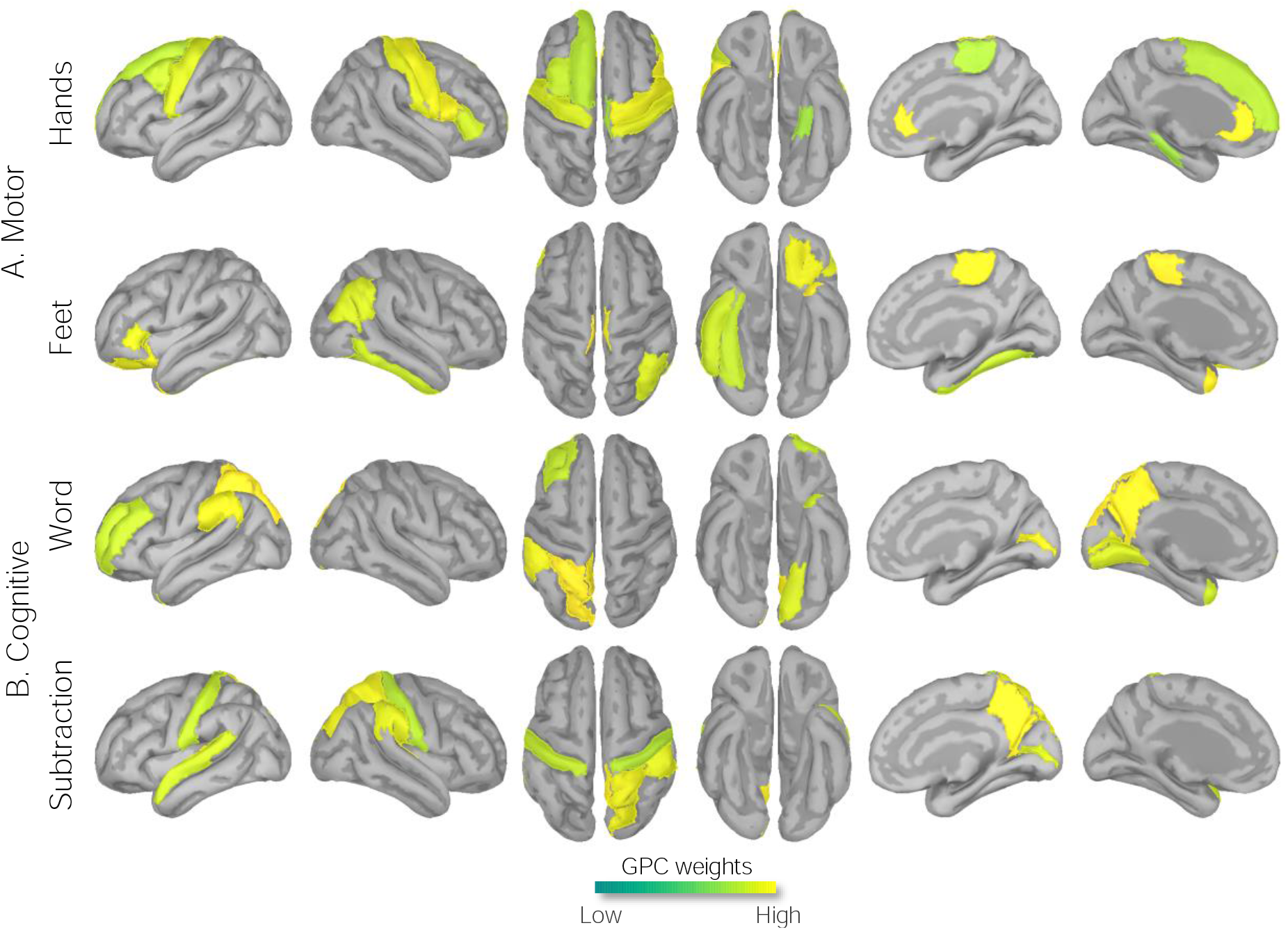
Weight (per region, Dk atlas) maps modeled by the GPC classification of BCI tasks. GPC classification weight maps of HANDS-FEET-WORD-Sub tasks are shown. Maps are consistent with regions reported in Table. 4. Rendered weights are displayed on a surface template (MNI-152) with dark representing sulci and gray representing gyri. Suprathresholded values with values greater than half maximum are shown.

**Table 4:** ROI involved in the GPC classification weight maps of BCI imagery tasks. Regions contributed (in percent) to the GPC classification of HANDS vs. FEET vs. WORD vs. Sub imagery tasks are reported. ROIs with percentage values greater than half-maximum (arbitrary) are reported.

| **HANDS, ROI** | **Weight (%)** | **FEET, ROI** | **Weight (%)** | **WORD, ROI** | **Weight (%)** | **SUB, ROI** | **Weight (%)** |
| --- | --- | --- | --- | --- | --- | --- | --- |
| rostral anterior cingulate R | 5.42 | paracentral L | 5.43 | superior parietal L | 5.92 | transverse temporal L | 6.34 |
| rostral anterior cingulate L | 5.29 | temporal pole L | 5.04 | precuneus L | 5.62 | precuneus R | 5.44 |
| parsopercularis R | 5.13 | paracentral R | 4.61 | pericalcarine R | 5.26 | transversetemporal R | 4.97 |
| precentral R | 4.69 | lateral orbitofrontal L | 4.41 | supramarginal L | 5.01 | superior parietal R | 4.86 |
| postcentral R | 4.59 | pars triangularis L | 4.01 | pericalcarine L | 4.91 | supramarginal R | 4.69 |
| precentral L | 4.58 | inferior parietal R | 3.84 | lingual L | 4.74 | pericalcarine R | 4.15 |
| pars triangularis R | 4.43 | inferior temporal R | 3.77 | temporal pole L | 4.23 | superior temporal L | 4.14 |
| caudal middle frontal L | 4.35 | fusiform R | 3.76 | rostral middle frontal L | 4.21 | postcentral L | 4.06 |
| paracentral R | 4.3 |  |  | rostral anterior cingulate R | 4.11 | postcentral R | 4.0 |
| parahipocampal L | 4.27 |  |  | paracentral R | 3.99 | isthmuscingulate R | 3.97 |

### 3.3. Correlation analysis

Correlations of individual beta-decrement maps and templates defined by the class mean (averaged across tasks and sessions) ranged between 0.53 and 0.65 (Fig. 8 B), with subjects 5 and 10 showing the highest correlations of *r* = 0.64 and 0.65, respectively. Across the two task sessions, participant 3 showed the most improved correlations (+11%) from sessions 1 to 2, but this improvement was not always the case (not seen in 5/14 subjects). The correlation between a subject’s beta-decrement map for a task and the class-mean template averaged across all fours tasks were *r* = 0.56 and 0.58, respectively, for sessions 1 and 2. A high ratio of correlations to the class-mean template, averaged over tasks, between sessions 1 and 2 (mean = 0.93, SD = 0.04) suggests consistency of cortical engagement across sessions. The correlation to class-mean showed significant positive correlations between sessions for the the WORD and HAND tasks (*r* = 0.57, p = 0.03 and 0.56, p = 0.03, respectively), whereas the SUB and FEET tasks showed non-significant correlations of *r* = 0.18 and 0.08, respectively. This suggests that the beta-decrement maps for these latter two tasks may be more unstable. Direct correlations of the beta-decrement maps across sessions for individual subjects (Fig. 8 G) appear to support this inference: they show the highest values for HAND and WORD task conditions (*r* = 0.52 and 57, respectively), and the lowest for the FEET task (*r* = 0.43). Fig. 8 summarizes these findings.

**Figure 8:**
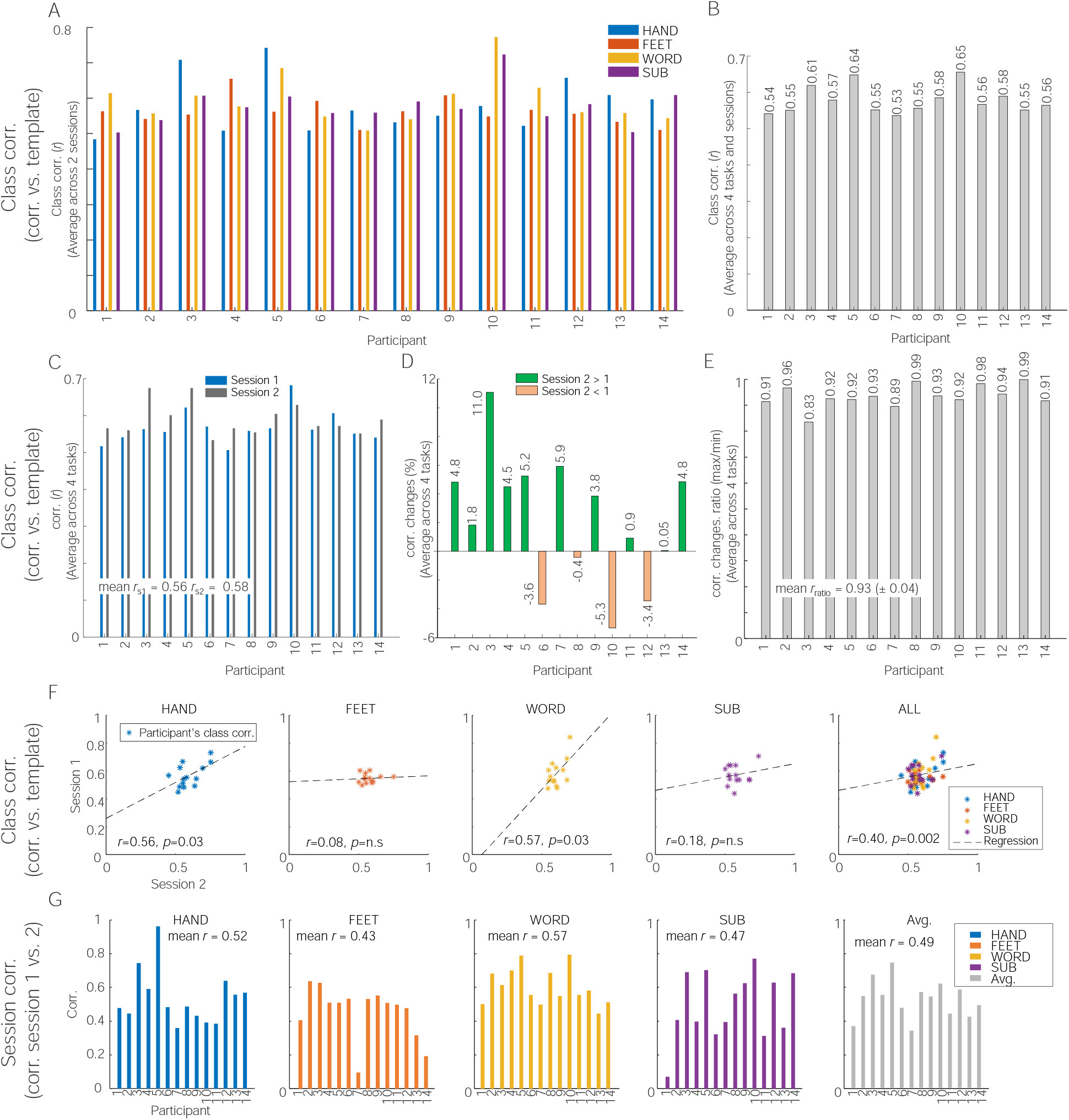
Correlation analysis of beta-decrements in BCI tasks. A Spearman correlation of beta-decrements of (A) four different imagery tasks (B) completed by 14 participants during (C) two sessions of a BCI experiment. (D) class correlation changes between sessions 1 and 2. The green and orange bars indicate increased and decreased correlation values for session 2 compared with session 1 of the task. (E) Correlation ratio of beta-decrements. The correlation ratio was computed by dividing the maximum over the minimum of individual class correlations between two sessions (F) scatter plots of class correlations of four BCI tasks. Linear Spearman correlations between sessions 1 and 2 are also reported, and a polynomial curve fitting line is overlaid. (G) Session-correlations, the correlation between sessions 1 and 2, of 4 BCI task conditions. Average session correlations are also shown and reported.

## 4. Discussion

This study set out to characterize and localize the neural correlates of motor imagery and mentally executed cognitive tasks in a group of healthy adult participants who completed an experimental paradigm that was designed for BCI research. We localized task-related cortical engagement from beta power decrements (Youssofzadeh et al., 2020) during motor imagery of the hands and feet, mental arithmetic (subtraction), and silent word-generation. Our source-space analyses at the group level revealed bilateral beta-desynchrony effects during motor imagery, notably in the motor region (pre-, post-, and para-central), the anterior portions of the cingulate gyri, middle-frontal gyri, and parts of the temporal neocortex. Group-level beta-decrements in the source space during the two cognitive tasks revealed left-lateralized effects in the parietal, temporal, and (inferior) prefrontal cortical regions.

Specifically, cortical engagement during motor imagery of the hands revealed bilateral beta power decrements in the supplementary motor area (SMA) and the precentral/primary motor cortex (BA 4), as well as anterior cingulate gyri (caudal and rostral) and prefrontal (caudal and medial orbitofrontal and frontal pole) regions (Figs. 4A and 5A and Table 1). The SMA, precentral, and postcentral gyri have both motor and sensory representations related to both upper and lower limbs (Jenkinson and Brown, 2011; Szameitat et al., 2007). The SMA is believed to support aspects of movement planning, including action preparation and the organization of movement sequences, and appears to have a role in the perception of stimuli that are potential targets of motor acts, and has been found to engage during cued movements (Bonini et al., 2014; Hardwick et al., 2013; Nachev et al., 2008; Lima et al., 2016; Bonini et al., 2014; Matsuzaka et al., 1992). The prefrontal cortex areas integrate information from the body and the environment and participate in higher-order gait control (Van der Meulen et al., 2014; Maidan et al., 2016). In particular, the medial frontopolar prefrontal cortex (BA 10) is involved in the integration of spatial and motor components of working memory during imagery and haptic exploration of spatial layouts, guiding motor preparatory processes (Kaas et al., 2007). We also found strong activation in the (caudal and rostral) anterior cingulate cortex (ACC) regions. The ACC (BA 32) lies in a unique position in the brain with connections to both the “emotional” limbic system and the “cognitive” prefrontal cortex (Bush et al., 2000). The ACC regions were shown to be engaged in a BCI motor imagery task requiring focused visual attention (Pfurtscheller et al., 2006; Luu and Posner, 2003). The ACC is also believed to play an important role in attentional control through bidirectional interactions with primary sensory areas (Crottaz-Herbette and Menon, 2006; Kim et al., 2016).

Group-level source analysis during motor imagery of the feet showed prominent beta power decrements in the paracentral/SMA and prefrontal (lateral orbitofrontal) regions; it also showed bilateral engagement of temporal regions (insula and superior temporal sulcus). The paracentral, precentral, and postcentral gyri also feature motor and sensory functions related to the lower limbs (Jenkinson and Brown, 2011). The paracentral lobules have the motor and sensory representations of the contralateral lower extremities, with sensory representations in its posterior portion (in the parietal lobe). Activation in paracentral/SMA has been found in studies of actual walking, using single-photon emission computed tomography and near-infrared spectroscopy (Fukuyama et al., 1997; Miyai et al., 2001). The superior temporal sulcus is a voice-selective area in the human auditory cortex and a source of sensory encoding, associated with motor output through the superior parietal-temporal area (Belin et al., 2018).

The anterior insula has been shown to be involved in mental navigation along memorized routes; it also supports the feeling of agency, awareness of body parts, and self-awareness (Van der Meulen et al., 2014; Ghaem et al., 1997; Craig, 2009). The group-level cortical engagements that we find during motor imagery of the feet are generally consistent with the results of the previous fMRI studies on gait imagery and limb movement imagery, supporting engagements in the primary and supplementary motor cortices and bilateral parietal and frontal areas (la Fougère et al., 2010; Van der Meulen et al., 2014; Bakker et al., 2007, 2008; Cojan et al., 2009; Wang et al., 2008).

Findings for the word generation and subtraction tasks showed strong left-hemispheric beta-decrement effects in the temporal (superior temporal gyrus), parietal (supramarginal gyrus), and (inferior) prefrontal gyri (Figs. 4B and 1B), regions known to be involved in cognitive (language) and comprehension (arithmetic) processing (Binder and Desai, 2011; Patterson et al., 2007; Arsalidou et al., 2018; Arsalidou and Taylor, 2011; Koyama et al., 2017). The supramarginal gyrus is an anatomical subdivision of the inferior parietal lobule, a heterogeneous brain region involved in the interpretation of both sensory and language information (Dehaene et al., 2004; Corbetta and Shulman, 2002; Cabeza et al., 2008). Both the left and right SMG regions are engaged in phonological (word) processing while the left SMG is more engaged in the semantic processing of lexical items (Oberhuber et al., 2016; Hartwigsen et al., 2010). Moreover, both Cognitive tasks revealed desynchrony effects in the IFG and postcentral (premotor) areas. The inferior parietal lobule is anatomically connected to ventral premotor areas, and the caudal inferior parietal gyrus is connected to the IFG regions (Petrides and Pandya, 2009; Caspers et al., 2013). Unlike the Word (linguistic) task, the Subtraction task led to greater beta power decrease effects in the left superior parietal lobule (SPL), whereas the Word condition task showed greater beta-power decrease effects in the left temporal regions, the inferior temporal gyrus (ITG), and the middle temporal gyrus (MTG). In general, mathematical calculations strongly engage the working memory, and the superior parietal lobule is critically important for the manipulation and rearrangement of information in the working memory processes (Koenigs et al., 2009; Roitman et al., 2012; Bemis and Pylkkänen, 2013). The ITG and its neighboring region, MTG, provide access to lexical-semantic representations during concept retrieval processes (Schuhmann et al., 2012; Hickok and Poeppel, 2007; Binder et al., 2009).

Our results from the pattern classification analyses support the use of beta-decrement mapping as a suitable approach for identifying cortical engagement related to BCI tasks from MEG recordings. Specifically, the SVM classification analysis of the BCI imagery task completion provided a reasonably high classification accuracy for two-way classification problems of HANDS vs. FEET (74%) and WORD-generation vs. SUBtraction (68%). Also, the GPC four-way classification accuracy of 60%, without any further dimension-reduction is reasonable given the complexity of combined task imagery responses. For our machine-learning pattern classification analyses, we utilized simple linear kernel values to train and classify four complex BCI problems. Generally, kernel methods are extremely useful for fast and efficient analyses. Kernels allow one to perform the learning using the kernel matrix, leading to an efficient computation instead of working with the whole data matrix, which avoids intensive computations. In addition to the computational advantages, kernels enable the solution of ill-conditioned problems and therefore avoid overfitting (Shawe-Taylor and Cristianini, 2004). The application of kernel methods to neuroimaging problems has been growing (Youssofzadeh et al., 2017; Schrouff et al., 2015; Chu et al., 2011), and maybe a potential candidate for real-time BCI applications.

The correlation analysis of beta-decrement effects reveals the consistency of responses between the two sessions in individual subjects, which may be related in part to the level of engagement with task imagery. For instance, higher class correlations were obtained for two participants, 10 and 5, which may suggest that these participants likely had better overall performance than others. This should ideally have been supported by behavioral data, and the lack of such measurements in the experimental paradigms that were employed is a key limitation of our study. This should be addressed in the design of future BCI paradigms (Klepp et al., 2015). Our correlation analysis indicated greater consistency of correlation to the group-mean for the HAND and WORD imagery conditions (*r* = 0.57 and 0.56, respectively) compared to the SUB and FEET tasks (*r* = 0.18 and 0.08, respectively), as shown in Fig. 8F. These findings are consistent with the lower (55%) binary classification accuracy of FEET vs. SUB and a high (85%) classification accuracy of HAND vs. WORD contrasts, as reported in Table 2, as well as the lower average session-to-session correlation of cortical-engagement maps for these two conditions (Fig. 8G) suggesting that mapping of mental subtraction and motor imagery involving the feet may be significantly less reliable than word generation and hand movement imagery tasks using beta-decrement effects.

Recent years have seen the increasing use of neuroimaging methods in the context of BCI research. Several studies have investigated the feasibility of MEG in real-time neurofeedback experiments (Mellinger et al., 2007; Buch et al., 2008; Gerven et al., 2009; Florin et al., 2014; Boe et al., 2014; Foldes et al., 2015; Okazaki et al., 2015; Fukuma et al., 2015, 2016). These studies have used both sensor and source-level MEG signals to provide feedback aimed at modulating specific brain activities. Indeed, the high temporal and spatial resolution of MEG makes it inherently suitable for real-time applications. To examine the sources of task-related cortical engagement, most BCI studies rely on the early P300 or N400 ERP components. However, brain source activities of late components (e.g., P600) are usually not phased-locked to an event and, therefore, cannot be extracted by linear methods such as averaging; they, however, may be detected by power changes in the frequency domain (David et al., 2006; Pfurtscheller and Andrew, 1999). ERP source estimates are limited to slow responses due to the effects of signal-averaging in the time domain. By contrast, induced power changes in the beta or gamma bands can serve as signatures of task-related cortical engagement (Crone et al., 2006; Singh et al., 2002; Eulitz et al., 1996). Beta-band power decrements have been used for classification purposes and clinical rehabilitation, e.g., characterization of motor or cognitive impairments as a therapeutic marker for patients recovering from a cerebral stroke (Buch et al., 2008) or helping robot-assisted gait training of patients with Parkinson’s disease or motor disability (Gwin et al., 2011; Severens et al., 2012; Knaepen et al., 2015). The beta-decrement approach can be applied to not only MEG and low-cost EEG systems but also the relatively new technology of optically-pumped magnetometers that offer good signal-to-noise ratio, spatial resolution, and portability for measuring BCI task activities (Knappe et al., 2012; Boto et al., 2018).

In summary, our results demonstrate the feasibility of using oscillatory dynamics of MEG signals, particularly beta-band power decrements, for localizing cortical engagement during a set of tasks designed for BCI research that did not involve dynamic sensory stimuli or overt behavioral responses. These motor imagery and mentally executed cognitive tasks (Rathee et al., 2021) engaged task-specific networks of brain regions that were largely consistent with prior neuroimaging studies of similar tasks in healthy adults and known neuroanatomy. Our results lend further support to the idea that task-related beta-band power decrements are closely associated with neural engagement in the cerebral cortex (Pfurtscheller and Lopes da Silva, 1999; Engel and Fries, 2010) and may be suitable for MEG BCI applications. BCI techniques have been used successfully in cognitive and motor training, leading to improvements in the performance of athletes, musicians, highly skilled manual technicians such as surgeons (Toth et al., 2020; Meister et al., 2004; Rogers, 2006), as well as those with a severe physical disability or motor impairment (Machado et al., 2010). Our results suggest that BCI applications for communication and rehabilitation may potentially benefit from the use of beta-band power decrements for localizing task-related cortical engagement.

## 5. Code and data availability

MEG data were analyzed using a combination of FieldTrip v20190419 (fieldtriptoolbox.org) and Brain-storm v060320 (neuroimage.usc.edu/brainstorm) toolbox in MATLAB2019a (The Mathworks, Inc.). Custom Matlab batch scripts including a Brainstorm implementation of non-overlapping temporal windows DICS beamformer source analysis is available at, github.com/vyoussofzadeh/BCI-beta-desynchrony_2021. All machine learning modeling steps were performed using PRoNTo v2 (mlnl.cs.ucl.ac.uk/pronto).

## 6. Acknowledgments

This work is in part supported by the UKIERI Thematic Partnership project, ‘Advancing MEG based Brain-Computer Interface Supported Upper Limb Post-Stroke Rehabilitation (DST-UKIERI-2016-17-0128), and the Northern Ireland Functional Brain Mapping Facility project (1303/101154803), funded by InvestNI and Ulster University.

The authors declare no conflict of interest.

## 7. Author contributions

V.Y and S.R conceived the idea and developed the methods. G.P. oversaw data acquisition. S.R organized the data. V.Y. performed the visualization and analysis. V.Y., S.R, and G.P. validated the analysis. V.Y. and S.R. drafted the manuscript. All authors participated in critically reviewing and revising the manuscript.

